# Predicting gender from structural and functional connectomes via brain and population graph neural networks

**DOI:** 10.1101/2023.11.01.565175

**Authors:** Yinan He, Yi Hao Chan, Jagath C. Rajapakse

## Abstract

Gender differences in terms of structural and functional organization of the human brain have been extensively studied, but existing works have mostly been limited to single modalities. In this paper, we propose a graph attention network architecture (BrainGAT) that uses informative subject-level features extracted from multimodal brain graphs to construct a population graph for gender classification. We show that while the extracted subject-level features can be directly used for classification, using these graph embeddings to construct a population graph further improves model performance. On the gender classification task, BrainGAT outperforms baseline models and existing multimodal modeling approaches, achieving an accuracy of 83.13% on the Human Connectome Project dataset. Salient connections highlighted by BrainGAT include connections between the inferior parietal and dorsolateral prefrontal areas of the cortex for females, while connections within the posterior cingulate cortex are highly salient for males. In sum, BrainGAT enables multimodal data to be modelled via population graphs in a parameter-efficient way.

**Clinical Relevance:** Several neurological conditions exhibit significant differences across genders and multimodal studies on these diseases are increasingly prevalent. This work highlights gender differences of multimodal connectomes in neurotypical settings. These insights could help to separate multimodal disease biomarkers from fundamental gender differences.

## I. Introduction

Neuroimaging methods are widely used to study the human brain and much attention has been devoted to identifying the neural basis of behavioural differences across genders. Elucidating the differences between the male and female brain is a key problem in neuroscience due to its impact on any downstream analysis tasks such as disease biomarker discovery. For instance, autism spectrum disorder is more prevalent in boys [1], while major depressive disorder is more common in females [2]. Whether this phenomenon is due to fundamental gender differences is unclear as the notion that sexual dimorphism even exists still remains controversial [3]. Nevertheless, it is clear that some differences exist across genders and more work is needed to fully understand their significance, especially in multimodal settings.

One key limitation in most existing studies is their focus on a single neuroimaging modality. Biological systems like the brain are more completely described by combining structural and functional analysis. For example, structural MRI (sMRI) captures anatomical details while functional MRI (fMRI) captures the dynamics of brain activity via measuring changes in blood oxygenation. Studies using a single modality neglect cross-modal interaction effects. However, the scarcity and complexity of neuroimaging data makes multimodal analysis challenging as models tend to overfit on such datasets.

Existing multimodal neuroimaging analysis [4], [5] typically combines diffusion tensor imaging (DTI) and fMRI, which can be represented as structural connectivity (SC) and functional connectivity (FC) matrices. Recent works on training deep learning models on such connectome datasets have converged towards the use of graph neural networks (GNN). GNN is an intuitive fit to connectome datasets as these data are fundamentally graphs. It alleviates the overfitting problem by significantly reducing the number of model parameters, relative to alternatives such as vanilla deep neural networks [6] or convolutional neural networks customized for connectivity matrices [7].

There are three main approaches of constructing graphs from connectome data. Firstly, brain graphs represent each subject as separate graphs and capture intra-subject correlation between regions of interest (ROIs) [4]. Secondly, population graphs represent a group of subjects as a single graph, thus modelling inter-subject correlation. [8] It uses similarity between subjects as the graph edges and vectorized FC matrices (which tends to be very high dimensional) as the subject’s feature vector. This approach is not scalable to multiple modalities especially when feature selection is not desirable (e.g., in biomarker discovery, feature importance needs to be assessed across all features). Lastly, a recent work [9] proposed a method that considers both brain and population graph, but is limited to a single modality.

In this work, we propose BrainGAT, a novel GNN architecture that incorporates both brain and population graphs to perform gender classification from DTI and fMRI datasets. Inspired by [8], our method improves it by enhancing the subject feature quality of the population graph. Unlike existing work, BrainGAT enables both SC and FC matrices to be used simultaneously, allowing cross-modal interactions to be considered. Our method outperformed baselines approaches as well as existing multimodal modelling [4] and dual graph [9] approaches. Ablation studies reaffirm the value of combining brain graphs and population graphs in both single modality and multimodal settings. BrainGAT also provides a way to identify salient connections in the male and female brain that considers the influence of SC on FC. The generated insights are different from existing studies performed on single modalities, potentially revealing novel findings that should be further verified in future multimodal studies.

## II. Method

We propose a two-stage early fusion method, as shown in Fig. 1, for classification based on two graph attention networks (GAT) [10]. In population graphs, the quality of subject features is crucial. Directly using high-dimensional single-modality data as subject features for population graph misses out on the cross-modal interaction effects. Therefore, it is desirable to encode a holistic representation of each subject’s connectome by using multiple modalities. This introduces a new design question of the best way to aggregate the multimodal information. Simply concatenating data from multiple modalities increases the dimensionality of the subject feature, resulting in severe overfitting. Our method uses a multimodal brain graph for subject representation learning, which incorporates information from multiple modalities.

**Figure 1.**
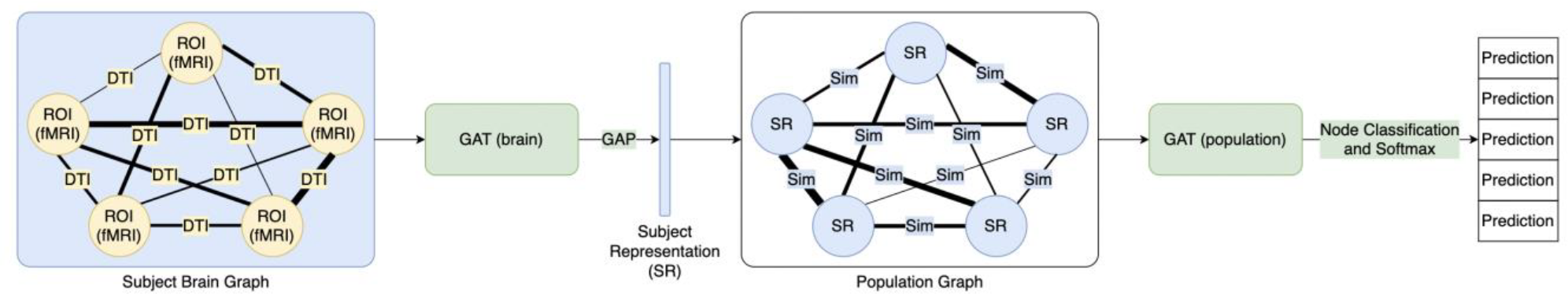
BrainGAT architecture

The stages of our method are:

1. Subject-level representation learning
2. Population graph construction and classification.

The subject representation learning stage performs early modality fusion and extract subject representations that are informative and have a reduced dimension. The extracted subject representation is used for population graph construction, where node classification is used for the final classification.

### A. Subject Representation Learning

We assume there are *n* ROIs for fMRI and DTI data. We construct a brain graph *G*^*B*^(*V*^*B*^, *E*^*B*^) using row vectors of FC matrix (fMRI) data as graph nodes and inter-node SC (DTI) strength as node edge weights. Node features are encoded in a feature matrix *X*^*B*^ ∈ ℝ^*n*×*n*^ whereby each 1 × *n* vector is the feature vector for an ROI. Edge weights are encoded in an adjacency matrix *A*^*B*^ ∈ ℝ^*n*×*n*^ whereby each element represents the SC strength between ROI pairs. We can use typical graph attention convolutions for feature learning. (1) defines the graph attention convolution operation on feature *X*, where Θ denotes the layer weights, *a* denotes the weights of a single-layer feedforward neural network for attention mechanism, *𝒩*(*i*) denotes the neighborhood of a node *i* in the graph.

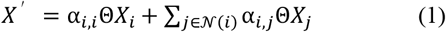

The attention coefficients α_i,j_ are computed as

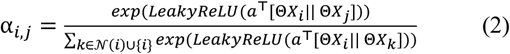

Global average pooling is applied after the last graph convolution to obtain the subject representation *S* ∈ ℝ^*n*×*d*^.

Let 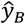 denote the predicted value and *y* be the ground truth. The subject representation extraction network is learned using the ground truth value of the subjects and cross-entropy loss on softmax activation output of the subject representation.

### B. Population Graph Construction and Classification

Using the learned subject representation, we construct a population graph *G*^*P*^(*V*^*P*^, *E*^*P*^). The node feature matrix is *S*, whereby each row is the learned representation of a particular subject. The edge weights *A*^*P*^ ∈ *R*^*n*×*n*^ represents the similarity between subject pairs. It is calculated using the similarity function defined in (3), where *corr*(·,·) computes the Pearson correlation coefficient for the two vectors. *i*_*VM*_ is the brain volumetric measures of subject *i*, which is the concatenated vector of total intracranial volume, grey matter, and white matter. These features are used to account for the effects of brain size differences on discovering male-female differences [3]. *i*_*SR*_ is the subject representation extracted from the brain graph. Graph convolutions stated in (1) are applied onto *S* with *A*^*P*^. Graph node classification is used to obtain individual subject prediction. Similar to subject representation learning, the population graph also learns using the ground truth value of the subjects and cross-entropy loss on softmax activation output.

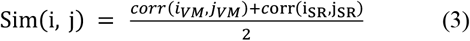

### C. Decoding

Decoding is done to obtain the saliency score for each feature (ROI). We performed decoding for both gender classes to identify features contributing the most to the classification decision. The saliency score indicates the importance of the feature to the classification task, such that a high saliency score (in terms of magnitude) implies a higher contribution from the feature. The saliency scores are obtained using the integrated gradients method [11].

Given a function *F* representing a deep neural network that maps input *x* to prediction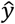, the saliency score *SS* for feature *i* of *x* is given by

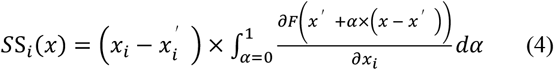

Where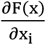 is the gradient of *F*(*x*) along the *i*^*t*h^ dimension and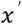 is the baseline of input *x*. The saliency score for an input feature *i* is the *i*^*t*h^ dimension of average saliency scores across all samples from the same class. In this study, saliency scores are aggregated across the class but it is possible to do the analysis at the level of individual patients by producing personalized disease biomarkers.

## III. EXPERIMENTS

### A. Dataset

The pre-processed HCP S900 version of the dataset was used in this study, involving 638 healthy young adults (283 male, 355 female) [12]. To generate the FC matrix, the Glasser atlas [13] was used to delineate 360 ROIs, and the time series of voxels within 2.5mm of the ROI were averaged. Pearson correlation coefficient is then computed on these 360 mean time series to obtain a symmetric functional connectivity matrix for each scan. Subjects in the HCP fMRI dataset often took multiple scans (ranging from 1 to 4). Each scan lasted for 15 minutes and subjects were told to keep their eyes open while keeping their gaze fixated on a screen with a white cross on a dark background. For the SC matrices, data processed by [14] were used. These matrices were also generated based on the Glasser atlas, thus producing SC matrices with 360 ROIs. These matrices were then used to build the brain graphs.

### B. Implementation Details

We tuned the model hyperparameters using 5-fold crossvalidation. These include: 1e-3 to 5e-3 for learning rate, 1e-4 to 5e-4 for weight decay in both training stages, 0.1 to 0.7 for dropout rate, and GNN layers including graph convolutional network (GCN), Chebyshev GCN and GAT. For both subject representation extraction network and population graph, we used a network architecture of 2 GAT convolution layers, each with 3 heads, exponential linear unit activation, and one linear layer. We used the Adam optimizer with weight decay of 3e-4 and a dropout rate of 0.5 for all layers. All networks are trained for 100 epochs with a batch size of 16 and 32 hidden channels. The subject features extracted from subject representation extraction networks have a dimension of 96. The experiments were done on an Nvidia P100 GPU.

### C. Experiment Results

We compared BrainGAT against logistic regression, dGLCN [9], and Joint-GCN [4]. For logistic regression, we applied both L1 and L2 regularization, with a ratio of 0.5 and a regularization strength of 1.0. The implementation of dGLCN and Joint-GCN are from the author’s published code. Our benchmarking results are presented in Table I. We show our method outperforms the second-best performing method by a large margin of 4.85%. The result for dGLCN on HCP dataset is unavailable as dGLCN requires the whole training set to be loaded as one batch. Considering the 360 × 360 SC and FC matrices, dGLCN would require over 62GB of GPU memory, which exceeds our GPU memory limit.

**TABLE I.**
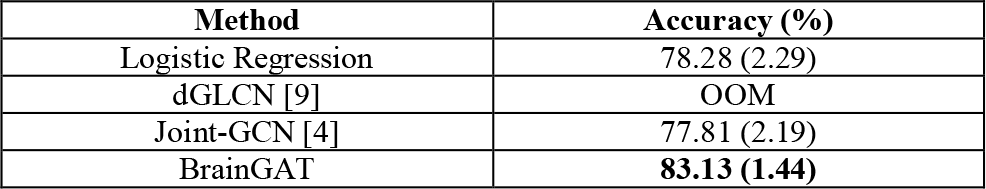
Comparisions with baselines and existing works.

### D. Ablation Studies

We conducted ablation studies on the design components of our method in Table II. Here, fMRI and DTI refers to using the two neuroimaging methods as modalities to construct the GNN. Brain refers to using brain graph and Population refers to using population graph for prediction. When only one modality is used for brain graph, it refers to using a single modality for both subject features and graph edges. When both fMRI and DTI are checked for brain graph, fMRI is used to construct the graph node vector and DTI is used as the graph edge weights. For methods using only population graph, the modality data is directly vectorized to form the subject feature. Our proposed method corresponds to the last row, where all design components are checked.

**TABLE II.** ablation studies for gender classification task.

| fMRI | DTI | Brain | Population | Accuracy (%) |
| --- | --- | --- | --- | --- |
| ✓ |  | ✓ |  | 79.69 (2.26) |
|  | ✓ | ✓ |  | 55.47 (0.00) |
| ✓ | ✓ | ✓ |  | 82.34 (1.53) |
| ✓ |  |  | ✓ | 57.19 (3.43) |
|  | ✓ |  | ✓ | 55.47 (0.00) |
| ✓ | ✓ |  | ✓ | 55.47 (0.00) |
| ✓ |  | ✓ | ✓ | 80.63 (1.81) |
| ✓ | ✓ | ✓ | ✓ | **83.13 (1.44)** |

Results from our ablation studies showed that using multiple modalities indeed improves the performance when using only brain graph or using both brain and population graph. This trend, however, does not hold when using only population graph. This could be because when using only population graph, the subject features are obtained by vectorizing the matrices and concatenating them when needed. In such a scenario, the dimensionality of the subject feature would be 129,240 for HCP dataset if two modalities are used. The high feature dimensionality aggravates overfitting, causing performance to deteriorate when multiple modalities are used for the population graph only baseline. We observe that using multiple modalities and using population graph improves the model performance, thus elucidating the importance of each design component.

### E. Visualisation of Learnt Representations

Subject feature quality is crucial to the performance of the population graph. Our ablation studies in Table II showed that using subject features extracted from the brain graph significantly increases the performance of the population graph as compared to using vectorized data.

We visualized the subject features using t-SNE [15] and the results corroborates our claim. As shown in Fig. 2, subject features extracted from brain graph are better separated by their class and form two clear clusters, whereas subject features directly obtained using concatenated vectorized SC and FC matrices show no clear clusters.

**Figure 2.**
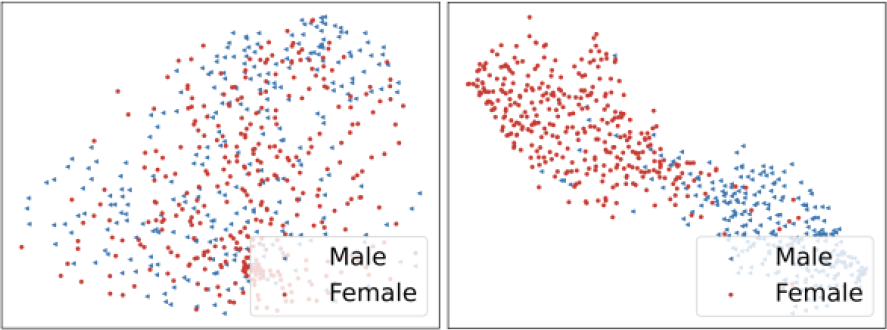
t-SNE plots of subject features using concatenated vectorized SC and FC matrices (left) and subject representation extracted from multimodal brain graphs (right).

### F. Decoding Results

Saliency scores were generated for every subject (only the subjects in the test split were used) and every feature. The baseline was defined as a vector of zeros and the scores were generated for each class separately (male and female). Since there were 5 folds in the cross-validation setup, 5 sets of scores can be generated. These scores are then averaged across all folds. Finally, the scores are averaged across subjects to generate class-wide saliency scores that show the features which are being focused on for the specified class.

Since the baseline is set as the zero vector, features that have high positive salience scores have much larger values than zero, while features with high negative salience scores have values much lower than zero. Amongst these most salient features, there will be some features shared across genders, but our key interest is on the differences. Table III and IV summarise the top (+ sign) and bottom (-sign) 0.005% of features that are different in females and males respectively.

**TABLE III.** Salient features for female subjects.

| Sign | ROI_1 | ROI_2 | Cortical regions involved |
| --- | --- | --- | --- |
| - | PFm_L | 9a_L | Inferior_Parietal, Dorsolateral_Prefrontal |
| - | PGi_R | 7m_R | Inferior_Parietal, Posterior_Cingulate |
| - | PGi_L | 9a_L | Inferior_Parietal, Dorsolateral_Prefrontal |
| - | LO1_R | LO2_R | MT+_Complex_and_Neighboring_Visual_Areas |
| + | PGi_R | v23ab_R | Inferior_Parietal, Posterior_Cingulate |
| + | PGi_R | 10d_L | Inferior_Parietal, Orbital_and_Polar_Frontal |

**TABLE IV.**
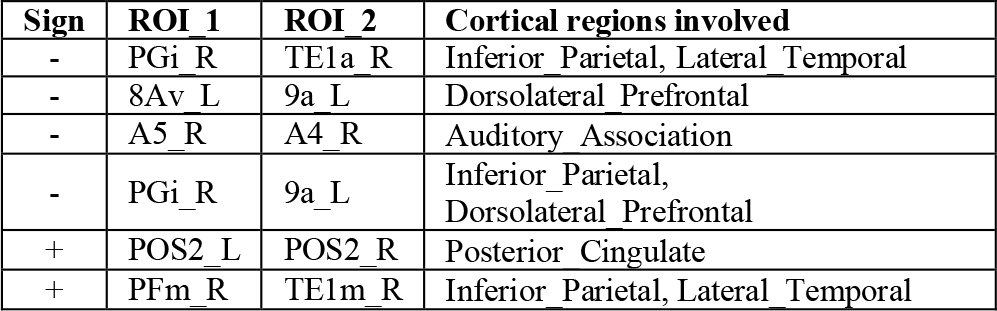
Salient features for male subjects.

| Sign | ROI_1 | ROI_2 | Cortical regions involved |
| --- | --- | --- | --- |
| - | PGi_R | TE1a_R | Inferior_Parietal, Lateral_Temporal |
| - | 8Av_L | 9a_L | Dorsolateral_Prefrontal |
| - | A5_R | A4_R | Auditory_Association |
| - | PGi_R | 9a_L | Inferior_Parietal, Dorsolateral_Prefrontal |
| + | POS2_L | POS2_R | Posterior_Cingulate |
| + | PFm_R | TE1m_R | Inferior_Parietal, Lateral_Temporal |

For females, multiple connections between the inferior parietal and dorsolateral prefrontal areas of the cortex had highly negative saliency scores, suggesting that these areas have relatively lower functional connectivity than others. For males, connections within the posterior cingulate as well as inferior parietal and lateral temporal cortical regions were found to have higher positive saliency scores. Regions such as inferior parietal and dorsolateral prefrontal areas of the cortex had highly negative saliency scores.

Overall, many connections identified in our study are unique. It could be possible that these regions are only elucidated when both SC and FC are considered, and they would not have been identified in studies that are limited to a single modality. Further research on gender differences using multimodal datasets will be needed to ascertain the robustness of these gender differences highlighted in our study.

## IV. CONCLUSION

BrainGAT provides a multimodal modelling approach that makes it possible to take in high-dimensional connectome features while still retaining the use of population graphs. Our method removes the need for feature selection and makes it possible to identify salient features – here, we identified key gender differences. Future work could extend this analysis to analyse gender differences in disease populations, explore the utility of intermediate fusion instead of the early fusion approach taken in this work, as well as to combine the two-stage training process into a single stage.

